# Scaling a context-aware genomic model for condition-specific edit design in plants

**DOI:** 10.64898/2026.10.23.761389

**Authors:** Paride Antinucci, Linfeng Wang, Jacob D. Moss, Dennis Schwartz, Dominic Hall

**Author notes:** Correspondence: Dominic Hall,.

## Abstract

Engineering agronomically relevant traits in crops often requires changing a gene’s activity in a specific tissue, developmental stage or environmental condition while preserving its roles in other contexts. Designing such selective changes requires predicting how cis-regulatory sequences are interpreted in different biological states, as the same edit can have different effects across contexts. Genomic models have made substantial progress in predicting gene expression from DNA sequence, yet sequence-only predictors do not receive an explicit description of biological states. We introduce CROP-1 (**C**ontext-aware **R**egulatory **O**utput **P**rediction model), which combines DNA sequence with transcription factor binding motifs and measured expression to predict gene activity as a function of both sequence and regulatory state. On a held-out test set spanning six plant species, CROP-1 achieves a median per-gene Pearson correlation across contexts of 0.831, compared with 0.486 for its parameter-matched sequence-only counterpart, and 57–246% higher median correlations than three genomic foundation models fully fine-tuned on the same task. Prediction accuracy also improves with scale: increasing model capacity reduces validation error by 48%, while expanding training data to include more diverse biological contexts reduces it by 58%. We then use gradient-based sequence optimisation to translate a desired gene expression pattern into candidate edits of its cis-regulatory sequence, targeting up- or down-regulation in one context while preserving expression in others. In soybean and tomato, edit designs with 1–20 nucleotide substitutions produce strong predicted target responses and much smaller average changes across non-target contexts. By connecting scalable learning of sequence–state relationships to selective edit design, CROP-1 provides a computational foundation for controlling where, when and under which conditions endogenous crop genes are active.

## Introduction

Crop improvement often requires changing gene activity in specific tissues, developmental stages or environmental conditions while preserving its function in other contexts. Many agronomic regulators affect several biological processes, so gene knockout or constitutive overexpression can couple a desirable trait to unwanted pleiotropic effects. Cis-regulatory editing offers a way to adjust when and where an endogenous gene is active while retaining the protein’s sequence and function. This approach has generated quantitative trait variation^1^, improved maize yield under drought^2^, separated competing rice yield components^3^ and conferred heat-stress resilience in tomato and rice by introducing heat-responsive regulatory elements^4^. Despite these advances, selecting regulatory edits for a new gene or crop remains difficult because the same sequence change can have different effects across biological contexts. A general design approach must therefore predict both the desired response in the target context and potential changes elsewhere, enabling candidate edits to be prioritised before experimental testing. Our aim is to develop a scalable model that learns how regulatory sequence and biological state jointly determine expression, and to use its predictions to guide the design of such edits in endogenous plant genes.

Sequence-to-function models provide a foundation for this approach by learning relationships between DNA and regulatory activity^5,6^. Enformer combines convolutional layers and self-attention to integrate long-range sequence information when predicting genomic tracks^7^. Borzoi extends this approach to higher-resolution RNA-seq coverage across gene bodies, enabling the analysis of transcription, splicing and polyadenylation within one model^8^. AlphaGenome further combines megabase-scale inputs with high-resolution predictions across expression and chromatin modalities^9^. These developments expand both the regulatory distances and the molecular processes that can be captured from sequence.

Self-supervised genomic pretraining complements these advances by learning from the much larger supply of unlabelled DNA. In plants, AgroNT and PlantCAD2 demonstrate how pretraining across diverse genomes can support regulatory prediction, variant interpretation and transfer to downstream tasks^10,11^. Nucleotide Transformer v3 connects multi-species sequence pretraining with functional supervision, supporting genomic-track prediction and controllable generation^12^. The BOTANIC-1 family extends plant genomic modelling through pretraining across 320 species and sequence contexts up to 128 kb, reporting state-of-the-art aggregate performance across 22 plant genomic tasks and strong fine-tuning results on regulatory prediction tasks^13^. These models provide increasingly capable sequence representations. However, these sequence-only predictors do not take an explicit description of biological state as input, leaving the regulatory context in which an edit acts implicit in their learned representations and task-specific outputs.

The need to represent biological state follows from how regulatory DNA is interpreted. Transcription factor (TF) binding motifs describe which sequences a factor can recognise^14^, while its abundance and activity help determine which interactions occur in a particular context. Plant promoter assays show that sequence features contribute differently across assay systems^15^, and enhancer mutagenesis reveals that interactions between regulatory elements change between light and dark conditions^16^. Context-aware models incorporate aspects of this variation as explicit inputs: GET combines sequence-derived features with chromatin accessibility^17^, Corgi uses regulator expression to modulate sequence features through feature-wise linear modulation^18^, and RegFM connects regulatory sequence representations to TF and chromatin-regulator activity through cross-attention^19^. Supplying context in this way allows a shared model to predict gene expression from the same DNA sequence across measured biological states, without requiring a separate output for each state. In plants, the broad coverage of transcriptomic resources makes TF expression a practical input, while binding-motif annotations allow species-specific TF repertoires to be represented within a shared predictor.

A shared predictor also creates an opportunity to learn from regulatory information distributed across species and experiments. Evolutionarily informed expression modelling and cross-species genomic pretraining support pooling information across taxa before refining a model for an individual crop^20,21^. How much can be gained from this approach depends on both model capacity and the scale and composition of the training data. Neural scaling studies describe systematic relationships between model size, data and predictive loss^22,23^, while data-selection studies show that informative examples can be more valuable than indiscriminate increases in dataset size^24^. This distinction matters for plant transcriptomics, where many experiments sample related tissues or regulatory states. Tissue aggregation can provide compact supervision, whereas individual experiments retain finer developmental and environmental variation. Comparing these alternatives can establish whether predictive gains depend primarily on more measurements, broader context coverage or greater model capacity.

For crop engineering, the purpose of these predictive gains is to guide useful regulatory changes. Large libraries of randomly synthesised promoters have enabled models to learn sequence–activity relationships^25,26^ and engineer expression in yeast^27^. Model-guided design has also produced cell-type-selective regulatory elements in mammals^28^ and tissue-specific enhancers in Drosophila^29^, demonstrating that biological specificity can itself be a design objective. Applying this principle to endogenous crop genes requires specifying both the contexts to alter and those to preserve, ideally through a small number of sequence changes. Such designs offer a practical connection to prime editing and its implementation in crops^30,31^. Explicit substitution budgets are also relevant to edit designs intended to meet European new genomic techniques (NGT) category 1 criteria, which constrain the number and size of genetic modifications alongside other eligibility requirements^32,33^.

Here we introduce CROP-1 (**C**ontext-aware **R**egulatory **O**utput **P**rediction model), which predicts gene expression by connecting DNA sequence features to TF context through cross-attention. A TF expression profile records the measured expression of selected TFs in a tissue or experiment; pairing these values with the corresponding DNA-binding motifs forms the TF context. This representation is intended to provide a compact yet granular description of regulatory state, capturing variation within and across tissues, developmental stages and conditions. Across six plant taxa, we assess the contribution of TF context against sequence-only and fully fine-tuned genomic foundation models, quantify improvements from model and data scaling, and test multi-species pretraining followed by taxon-specific fine-tuning (Fig. 1a). We then use gradient-based sequence optimisation to design sparse substitutions in soybean and tomato that increase or decrease predicted expression in a target tissue context while limiting changes in non-target contexts.

**Figure 1.**
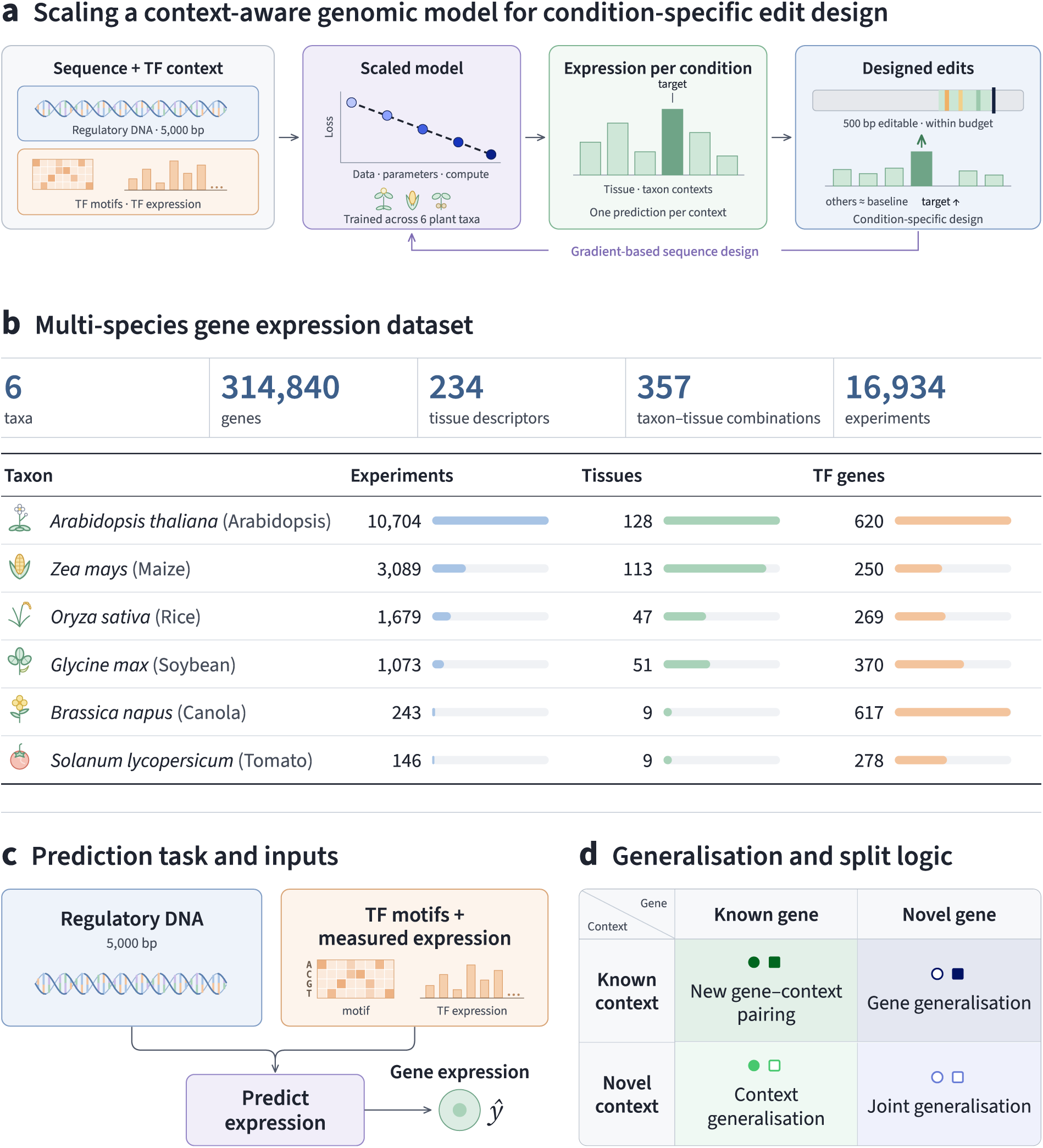
A framework for learning and designing context-specific gene expression in plants. **a**, Study workflow linking DNA and TF-context inputs to model scaling, benchmarking, multi-species transfer and sequence-edit design. **b**, Dataset inventory across six plant taxa. Experiment counts refer to TF expression profiles; genes are counted across the tissue-benchmark training and test sets. Tissue descriptors and taxon–tissue combinations are distinct measures of context coverage. **c**, A gene’s 5-kb sequence and a TF context jointly determine one expression prediction. **d**, Four test strata separate generalisation across genes, contexts and their combinations. Filled symbols indicate known entities and open symbols indicate novel entities. Novel contexts are taxon–tissue combinations, with TF measurements available as input; gene novelty is defined by gene identifiers.

## Results

### A shared expression predictor across six plant taxa

We first established a benchmark to test whether a single sequence-and-context model could predict gene expression across plant species and biological states. Using PlantExp-derived expression data^34^, we assembled a resource spanning *Arabidopsis thaliana*, *Brassica napus*, *Glycine max*, *Oryza sativa*, *Solanum lycopersicum* and *Zea mays*. It contains 314,840 genes, 234 tissue descriptors and 357 taxon–tissue combinations, supported by 16,934 experiment-level TF expression profiles (Fig. 1b). Each profile records the expression levels of the TFs measured in one experiment; these values are paired with the corresponding DNA-binding motifs to form the TF context supplied to the model.

To provide consistent supervision across this heterogeneous collection, we first constructed a tissue-aggregated benchmark. For each gene–taxon–tissue combination, we transformed experiment-level TPM values using the natural logarithm, log1p(TPM), then took their median across experiments; tissue-level TF expression profiles were aggregated in the same way. This produced approximately 14.5 million training observations and 0.9 million test observations. Each observation pairs a gene’s sequence with a tissue-level TF context and an expression target. We used this compact dataset for the model-scaling and sequence-model comparisons, before testing what could be gained by retaining individual experiments.

CROP-1 predicts one expression value from a 5,000-bp genomic sequence and the TF context of the queried tissue or experiment (Fig. 1c). The window spans 3 kb upstream and 2 kb downstream of the annotated 5′ gene boundary, used here as the transcription start site (TSS) reference, and is oriented in the direction of transcription. For each taxon, we selected TF genes with binding motifs annotated in PlantTFDB^35,36^ and matched them to the available expression measurements. Each TF contributes a binding-motif representation and its measured expression, and the number of TFs can vary across taxa. The same scalar-output predictor is shared across genes, species and contexts. This formulation lets the model predict expression for a given gene in different biological states by changing the supplied TF expression profile.

We evaluated generalisation along two axes: gene identity and taxon–tissue context. We stratified the test set—that is, divided it into groups according to exposure during training—to distinguish four settings: known genes in known contexts, known genes in novel contexts, novel genes in known contexts, and novel genes in novel contexts (Fig. 1d). All six taxa and all 234 tissue descriptors are represented somewhere in the training data, but selected taxon–tissue combinations are withheld. A novel context therefore pairs a represented taxon with a tissue for which that taxon provides no training targets.

Measured TF expression profiles are supplied at prediction time in every group. Novelty refers to the target supervision withheld during fitting: gene holdouts test transfer across sequence inputs, context holdouts test the use of TF expression profiles for withheld taxon–tissue pairings, and the joint holdout tests both together. These evaluations assess whether learned relationships between sequence and regulatory state extend beyond the supervised gene–context pairs, an important prerequisite for using the model to guide sequence design.

Two complementary metrics assess these predictions. Tissue-level Pearson correlation compares expression across genes within a tissue descriptor, whereas gene-level correlation compares expression across contexts for an individual gene. Unless stated otherwise, tissue-level correlations subtract the same gene-specific median from predictions and observations to focus on expression relative to each gene’s typical level; we examine the importance of this adjustment below. The benchmark thus tests both which genes respond within a context and how an individual gene responds across contexts, including combinations not encountered during training.

### Model scaling systematically improves expression prediction

We next asked whether increasing model capacity improves the accuracy of context-aware expression prediction and its ability to generalise. Across five configurations spanning approximately 4 million to 400 million parameters, error on the held-out validation set decreased monotonically with model size, accompanied by improved prediction on held-out genes and contexts.

CROP-1 combines a convolutional sequence backbone with TF cross-attention at multiple depths (Fig. 2a). Residual convolutional blocks extract sequence features, which supply attention queries; TF tokens combine motif features with context-specific expression and supply keys and values. The full set of motif–expression tokens constitutes the TF context. Separate learned attention modules at successive depths connect the same TF context to progressively more abstract sequence features. Pooling and a dense head produce the expression prediction. When the target gene is itself a TF, its expression value is withheld and its token excluded from attention, preventing direct recovery of the target from the context input.

**Figure 2.**
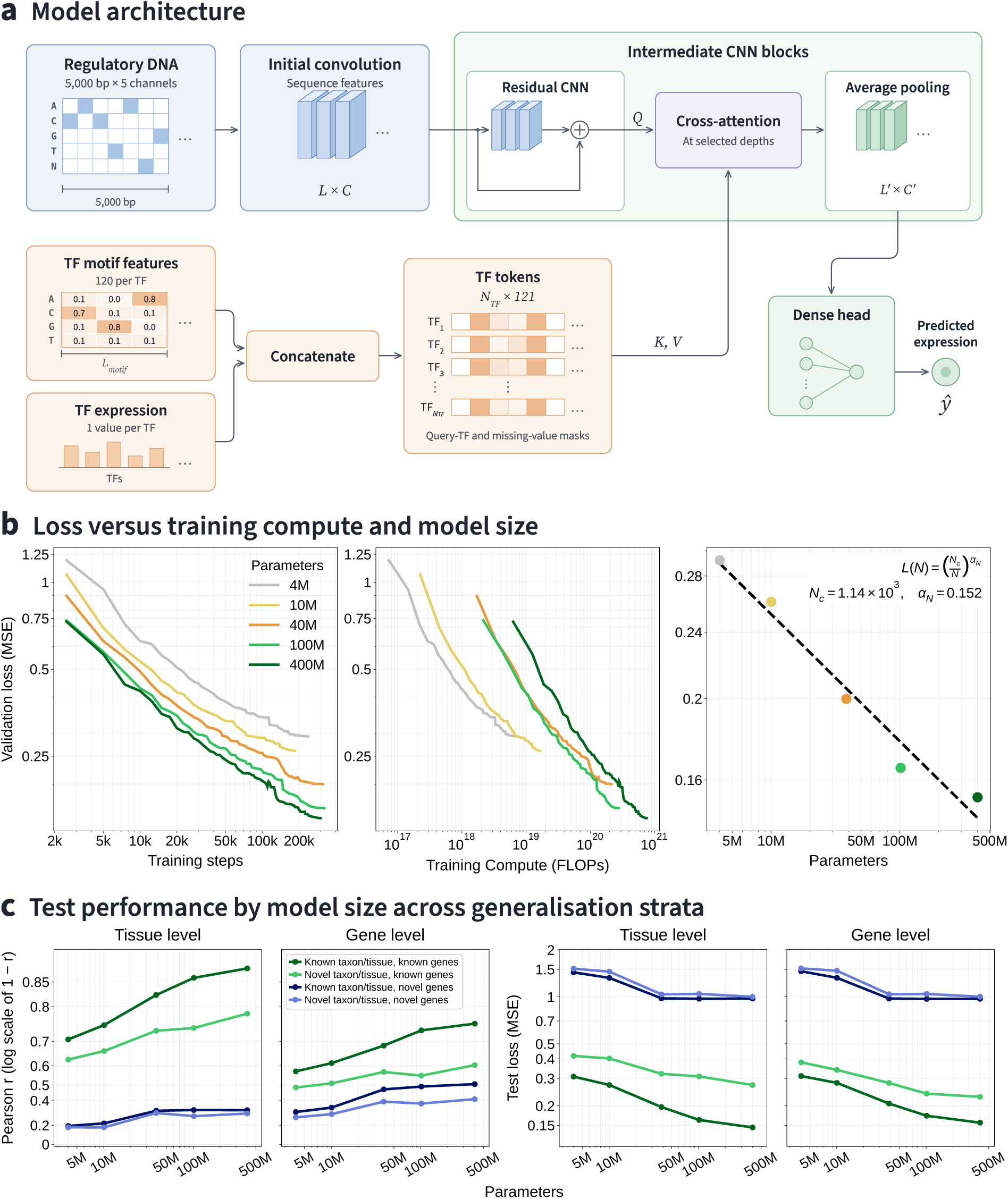
Architecture and scaling of CROP-1. **a**, Residual convolutional neural network (CNN) blocks extract sequence features, which interact with motif–expression TF tokens through cross-attention at selected depths. Pooling and a dense head produce the expression prediction. Total model parameter counts are reported in Table 4. **b**, Running minimum validation MSE against optimiser updates and estimated cumulative training FLOPs, followed by validation MSE at convergence against model size. The empirical fit is *L*(*N*) = (*N_c_*/*N*)^*α*^*^N^*, with *N_c_* = 1.14 × 10^3^ and *α_N_* = 0.152. Models are labelled by rounded parameter counts. **c**, Mean tissue- and gene-level Pearson correlation and MSE across the four generalisation strata. Correlation uses a − log_10_ (1 −*r*) display transformation, with tick labels in correlation units. Tissue-level correlations use predictions and observations centred by the same gene-specific median (Methods).

Increasing capacity nearly halved validation error. With all five models trained to convergence using a common optimiser schedule and global batch size, validation MSE decreased from 0.2923 at 4M parameters to 0.1525 at 400M, a 47.8% reduction (Fig. 2b). Loss decreased at every increase in size, including a further 7.7% reduction between 100M and 400M. The converged losses followed an empirical power law with parameter count, with exponent 0.152 and log-space R^2^ of 0.967. Training trajectories also showed decreasing loss with additional compute within each configuration.

On the test set, the gains extended to both evaluation axes and all four generalisation settings (Fig. 2c). For known genes in known contexts, mean tissue-level Pearson correlation rose from 0.707 to 0.872 between the smallest and largest models, while mean gene-level correlation rose from 0.574 to 0.756. For known genes in novel contexts, gene-level correlation increased from 0.485 to 0.604. Novel genes also benefited: tissue-level correlation in known contexts increased from 0.193 to approximately 0.33, and the largest model reached a mean gene-level correlation of 0.410 when both genes and contexts were novel.

Model capacity is therefore an effective scaling axis for context-aware gene expression prediction in plants. The improvements across both tissue- and gene-level metrics show that larger CROP-1 models better predict context-dependent expression, including responses beyond the genes and contexts used for training.

### TF context improves prediction beyond large pretrained sequence models

To determine whether explicit TF context adds predictive information beyond sequence-model capacity and genomic pretraining, we compared CROP-1 with a parameter-matched model without TF context and three genomic foundation models that we fully fine-tuned on the same expression task. CROP-1 achieved the highest mean Pearson correlationsacrossall eight gene- and tissue-level generalisation comparisons, with the clearest advantage in metrics that capture context-dependent expression (Fig. 3; Table 1).

**Figure 3.**
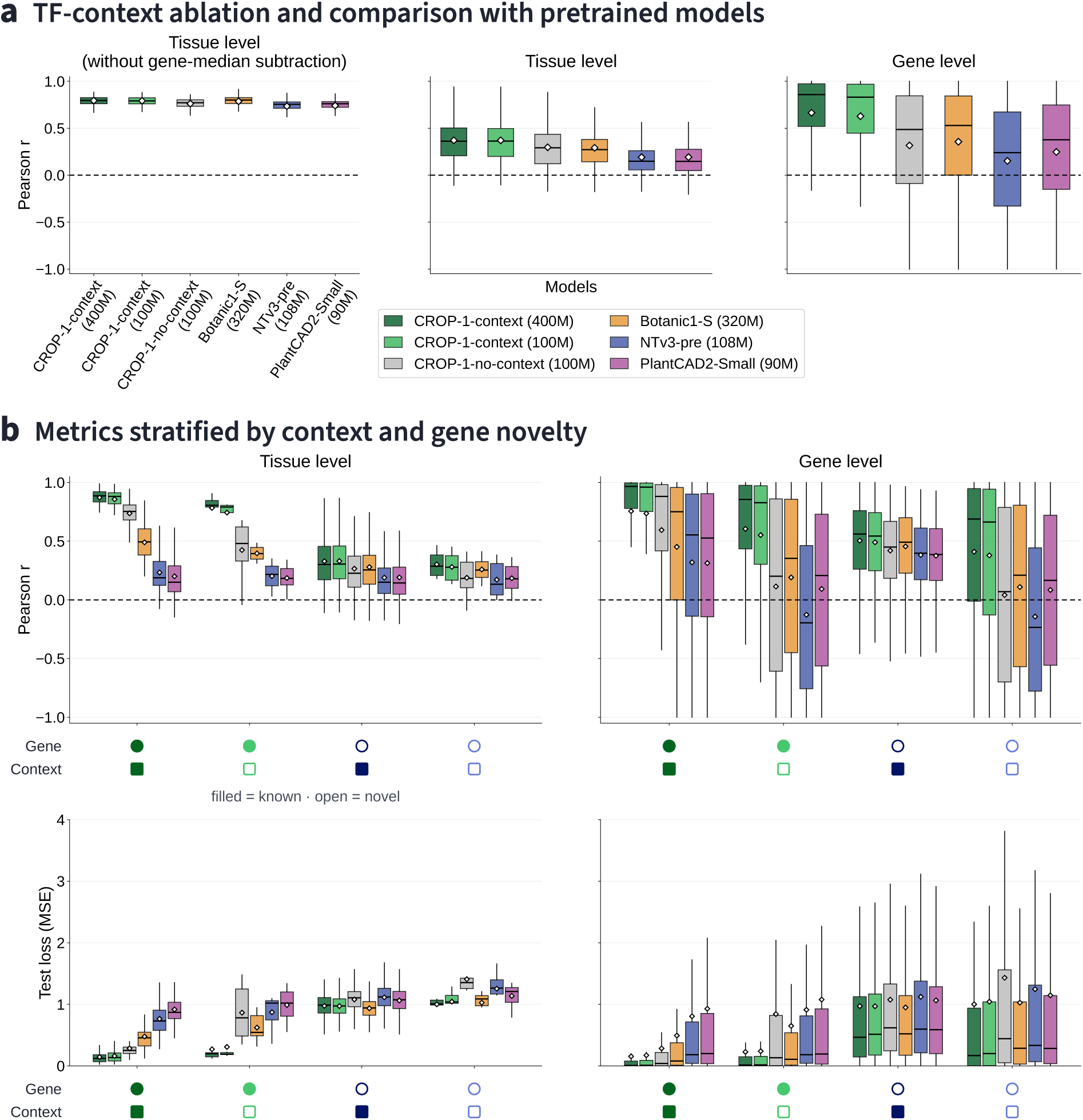
TF context improves overall and stratified expression prediction. Comparisons include CROP-1 at 400M and 100M parameters, the 100M CROP-1 model without TF context, and Botanic1-S, NTv3-pre and PlantCAD2-Small, each fully fine-tuned on the expression task. **a**, Non-stratified Pearson-r distributions after combining the test strata. Left, correlation across genes within each tissue before gene-median subtraction; middle, the corresponding tissue-level correlation after subtracting the same gene-specific median from predictions and observations; right, correlation across contexts within each gene. The reference median is calculated from tissue-aggregated training and test labels for evaluation only (Methods). Subtraction removes each gene’s typical expression level from the tissue-level comparison and leaves gene-level correlation and MSE unchanged. **b**, Pearson correlation (upper row) and MSE (lower row), stratified by gene and taxon–tissue context novelty, at tissue level (left) and gene level (right). Tissue-level correlations use gene-median subtraction. Filled and open symbols denote known and novel groups. Boxes show the interquartile range, black lines the median, white diamonds the arithmetic mean and whiskers the most extreme values within 1.5 interquartile ranges. Outlier markers are hidden without excluding those observations from summaries. Black dashed lines indicate Pearson r = 0. Numerical summaries are given in Table 1.

**Table 1.** Overall and stratified expression prediction with CROP-1 and fine-tuned genomic foundation models. **a**, Non-stratified Pearson correlations, calculated across genes within each tissue or across contexts within each gene, and summarised as mean (median). Tissue correlations are shown before and after subtracting the same gene-specific median from predictions and observations; gene-level correlation is unchanged by this subtraction. **b,c**, Stratified tissue- and gene-level performance, with each cell reporting mean Pearson r / mean MSE. Tissue-level correlations in **b** use gene-median-centred expression. Values are rounded to three decimal places; bold marks the best value for each summary statistic at the displayed precision. Model sizes are approximate.

**a. Non-stratified Pearson correlation: mean (median)**
| Model | Tissue $r$ | | Gene $r$ |
| --- | --- | --- | --- |
|  | Before subtraction | After subtraction |  |
| CROP-1 (400M) | **0.795** (0.795) | **0.371** (0.361) | **0.664** (0.858) |
| CROP-1 (100M) | 0.792 (0.792) | **0.371** (0.363) | 0.628 (0.831) |
| CROP-1-no-context (100M) | 0.762 (0.771) | 0.297 (0.292) | 0.316 (0.486) |
| Botanic1-S (320M) | 0.786 (**0.801**) | 0.292 (0.271) | 0.356 (0.529) |
| NTv3-pre (108M) | 0.735 (0.755) | 0.191 (0.149) | 0.152 (0.240) |
| PlantCAD2-Small (90M) | 0.741 (0.761) | 0.192 (0.147) | 0.247 (0.377) |

**b. Tissue-level performance: mean Pearson $r$ / mean MSE**
| Context / gene | CROP-1 (400M) | CROP-1 (100M) | CROP-1-no-context (100M) | Botanic1-S (320M) | NTv3-pre (108M) | PlantCAD2-Small (90M) |
| --- | --- | --- | --- | --- | --- | --- |
| Known / known | **0.872 / 0.145** | 0.857 / 0.162 | 0.737 / 0.285 | 0.490 / 0.479 | 0.233 / 0.765 | 0.200 / 0.918 |
| Novel / known | **0.783 / 0.272** | 0.743 / 0.309 | 0.425 / 0.865 | 0.396 / 0.623 | 0.203 / 0.872 | 0.186 / 0.989 |
| Known / novel | 0.329 / 0.975 | **0.331** / 0.972 | 0.266 / 1.080 | 0.279 / **0.936** | 0.190 / 1.114 | 0.191 / 1.063 |
| Novel / novel | **0.302 / 1.001** | 0.281 / 1.045 | 0.188 / 1.409 | 0.259 / 1.027 | 0.172 / 1.254 | 0.184 / 1.137 |

**c. Gene-level performance: mean Pearson $r$ / mean MSE**
| Context / gene | CROP-1 (400M) | CROP-1 (100M) | CROP-1-no-context (100M) | Botanic1-S (320M) | NTv3-pre (108M) | PlantCAD2-Small (90M) |
| --- | --- | --- | --- | --- | --- | --- |
| Known / known | **0.756 / 0.156** | 0.736 / 0.173 | 0.595 / 0.285 | 0.452 / 0.494 | 0.319 / 0.807 | 0.313 / 0.929 |
| Novel / known | **0.604 / 0.228** | 0.552 / 0.239 | 0.114 / 0.843 | 0.191 / 0.650 | -0.127 / 0.911 | 0.093 / 1.078 |
| Known / novel | **0.505** / 0.972 | 0.491 / 0.970 | 0.420 / 1.076 | 0.455 / **0.949** | 0.383 / 1.123 | 0.376 / 1.065 |
| Novel / novel | **0.410 / 1.002** | 0.379 / 1.044 | 0.041 / 1.432 | 0.109 / 1.028 | -0.142 / 1.246 | 0.084 / 1.147 |

The foundation-model baselines were Botanic1-S, NTv3-pre and PlantCAD2-Small^11,12,13^. Starting from their pretrained checkpoints, we fine-tuned each backbone in full together with a tissue-expression regression head. We compared these models with 100M CROP-1 and its 100M sequence-only counterpart. Available pretrained variants were selected to match the 100M model’s parameter count as closely as possible. The sequence-only models predict a fixed set of 234 tissue outputs, whereas CROP-1 receives TF context for the queried biological state. We also included 400M CROP-1 to connect this comparison with the capacity-scaling experiment above.

Gene-median subtraction was essential to distinguish prediction of context-associated variation from prediction of a gene’s typical expression level. Before subtraction, the six models achieved comparably high mean tissue-level correlations between 0.735 and 0.795; the upper value belonged to 400M CROP-1, while 100M CROP-1 and Botanic1-S reached 0.792 and 0.786, respectively (Fig. 3a). A model can predict the large baseline differences in expression between genes and thereby achieve high tissue-level correlation, even when it predicts little of each gene’s variation across contexts. We therefore subtracted each gene’s median expression from both its predicted and observed expression. Tissue-level correlation then measures agreement in expression relative to that gene-specific reference.

After this adjustment, the advantage of TF context was clear. On the non-stratified test set, both CROP-1 models achieved a mean tissue-level correlation of 0.371, compared with 0.297 for CROP-1 without TF context, 0.292 for Botanic1-S and approximately 0.19 for NTv3-pre and PlantCAD2-Small. The complementary gene-level analysis, which compares contexts within each gene, gave mean correlations of 0.628 for 100M CROP-1 and 0.664 for 400M CROP-1, versus 0.316 without TF context and 0.356 for Botanic1-S. Gene-median subtraction leaves within-gene correlation and MSE unchanged, because the same constant is subtracted from predictions and observations for a given gene.

The advantage persisted when gene and context novelty were examined separately (Fig. 3b). The CROP-1 family achieved the highest mean Pearson correlation in all eight stratum–axis comparisons, and 100M CROP-1 exceeded Botanic1-S in each comparison despite Botanic1-S having approximately three times as many parameters. For known genes in novel contexts, mean gene-level correlation was 0.552 at 100M and 0.604 at 400M, compared with 0.114 without TF context and 0.191 for Botanic1-S. When both genes and contexts were novel, 400M CROP-1 reached 0.410, compared with 0.041 and 0.109, respectively.

TF context also reduced prediction error: the 100M CROP-1 model had lower mean MSE than the parameter-matched sequence-only comparator in every stratum at both evaluation levels. The 400M model achieved the lowest mean MSE for known genes and for novel genes in novel contexts; Botanic1-S achieved the lowest mean MSE for novel genes in known contexts (Table 1).

In summary, explicit TF context improves prediction of expression changes across biological states beyond the tested sequence-only models, with the largest gains in gene-level correlation occurring in novel taxon–tissue contexts for both known and novel genes. Gene-median centring makes this benefit visible at tissue level by separating context-associated expression from the large baseline differences between genes.

### Tissue aggregation and context diversity enable efficient data scaling

We next examined how predictive performance scales with increasing coverage of experiment-level contexts, and how much of that performance can be achieved using tissue-aggregated training data. At a fixed model size of 100 million parameters, increasing experiment-level supervision substantially improved prediction, while the tissue-aggregated dataset retained much of the predictive signal with far fewer observations.

To expand the training data across distinct regulatory states, we clustered experiments within each taxon using correlation distances between TF expression profiles and selected one representative per cluster (Fig. 4a). Beyond tissue identity, these experiments cover a broad range of developmental stages and experimental conditions, providing variation in regulatory state within each tissue. Four thresholds yielded subsets labelled approximately 1%, 5%, 10% and 50%, followed by the complete training set (Fig. 4b). The resulting datasets ranged from 7.78 million to 568.30 million training observations (Table 2), increasing both expression coverage and the diversity of TF expression profiles supplied to the model.

**Figure 4.**
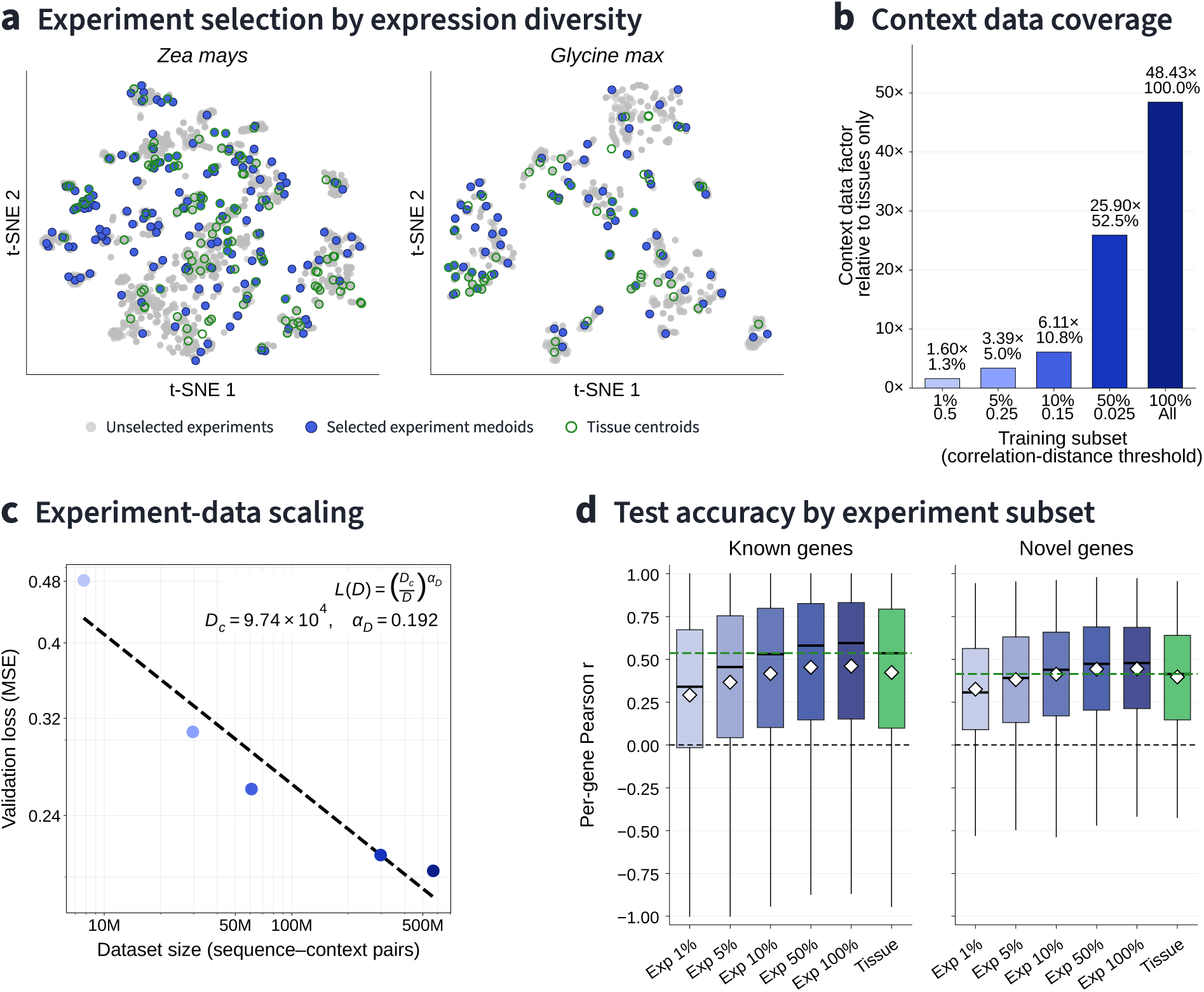
Context diversity and experiment-level data scaling. **a**, Independent t-SNE visualisations of TF expression profiles from maize and soybean, selected as two example species for display. Points show tissue-aggregated expression profiles and experiment representatives selected by clustering experiment-level TF expression profiles at a correlation-distance threshold of 0.25. Tissue points labelled “centroids” denote tissue-aggregated expression profiles; clustering is performed in TF expression space independently of the visualisation. **b**, Context-inventory ratios relative to 357 tissue-level TF expression profiles, calculated as (357 + selected experiment medoids)/357. These ratios describe context coverage; training uses the experiment observations listed in Table 2. **c**, Converged validation MSE against training-observation count, with empirical fit *L*(*D*) = (*D_c_*/*D*)^*α*^*^D^*, *D_c_* = 9.74 × 10^4^ and *α_D_* = 0.192. **d**, Distributions of per-gene Pearson correlation on the experiment test set for known and novel genes, comparing the five subsets with tissue-only training. Boxplot conventions follow Fig. 3. Green dashed lines indicate the tissue-trained model’s median.

**Table 2.** Experiment-level datasets used for scaling. Training counts exclude the fixed validation sample. Subset percentages are nominal labels for the clustering settings; the realised context and observation counts define each training dataset.

| Nominal subset | Correlation-distance threshold | Training contexts | Training observations | Validation MSE at convergence |
| --- | --- | --- | --- | --- |
| 1% | 0.500 | 210 | 7,779,788 | 0.481330 |
| 5% | 0.250 | 810 | 29,681,671 | 0.307417 |
| 10% | 0.150 | 1,693 | 60,985,693 | 0.259557 |
| 50% | 0.025 | 8,303 | 297,759,472 | 0.213508 |
| 100% | All eligible profiles | 15,899 | 568,303,917 | 0.203749 |

Validation MSE decreased by 57.7% across this range, from 0.4813 to 0.2037, with every configuration trained to convergence (Fig. 4c). The five converged losses followed an empirical power law with data exponent 0.192 and log-space R^2^ of 0.924. Gains continued through the largest expansion: using all training observations reduced validation MSE by a further 4.6% relative to the 50% subset.

The additional supervision also improved prediction on approximately 48 million experiment-level test observations, assigned using the same gene–taxon–tissue keys as the tissue-level train/test split (Fig. 4d). Between the smallest and largest training subsets, mean per-gene Pearson correlation increased from 0.291 to 0.461 for known genes and from 0.326 to 0.445 for novel genes. Broader experiment coverage therefore improved prediction of expression variation across experimental contexts for both gene groups.

Tissue aggregation nevertheless provided efficient supervision for this same task. To make predictions for individual experiments, we supplied the tissue-trained model with each experiment’s measured TF expression profile in place of the tissue-aggregated profile used during training. With roughly 40-fold fewer training observations than the complete experiment dataset, this model reached correlations of 0.423 for known genes and 0.397 for novel genes, and remained competitive in MSE. The strongest experiment-trained model improved correlations further, indicating that individual experiments contain useful context-dependent signal beyond the tissue summaries.

Tissue aggregation therefore captures substantial expression signal in a compact training set. By supplying TF expression measurements from an individual experiment, the tissue-trained model can predict gene expression in that experiment’s regulatory context. Training on a broader range of experiment-level contexts improves accuracy further, making data composition as well as observation count relevant to scaling.

### Multi-species pretraining improves taxon-specific prediction

We next tested whether multi-species pretraining improves prediction within a target species after taxon-specific fine-tuning. We compared three strategies at 100 million parameters: a single shared model trained jointly on the complete experiment data from all six taxa; separate models trained from scratch on each taxon; and the shared model fine-tuned in full on each taxon’s data.

The scratch-trained and fine-tuned models used the same target-taxon training data, differing in their starting parameters. All strategies were trained to convergence, so the comparison did not impose a common update limit that could leave models trained on different dataset sizes at different stages of optimisation. This addresses under-training as an explanation for the observed differences; multi-species pretraining still entails additional data exposure and compute.

Multi-species pretraining followed by taxon-specific fine-tuning gave the highest aggregate correlations for both known and novel genes (Fig. 5a). For known genes, mean Pearson correlation increased from 0.461 with the shared model to 0.537 with taxon-specific training from scratch and 0.560 after fine-tuning. Mean MSE decreased from 0.2993 to 0.2063 and 0.1766, respectively. Fine-tuning therefore reduced mean MSE by approximately 14% relative to a converged model trained on the target taxon alone.

**Figure 5.**
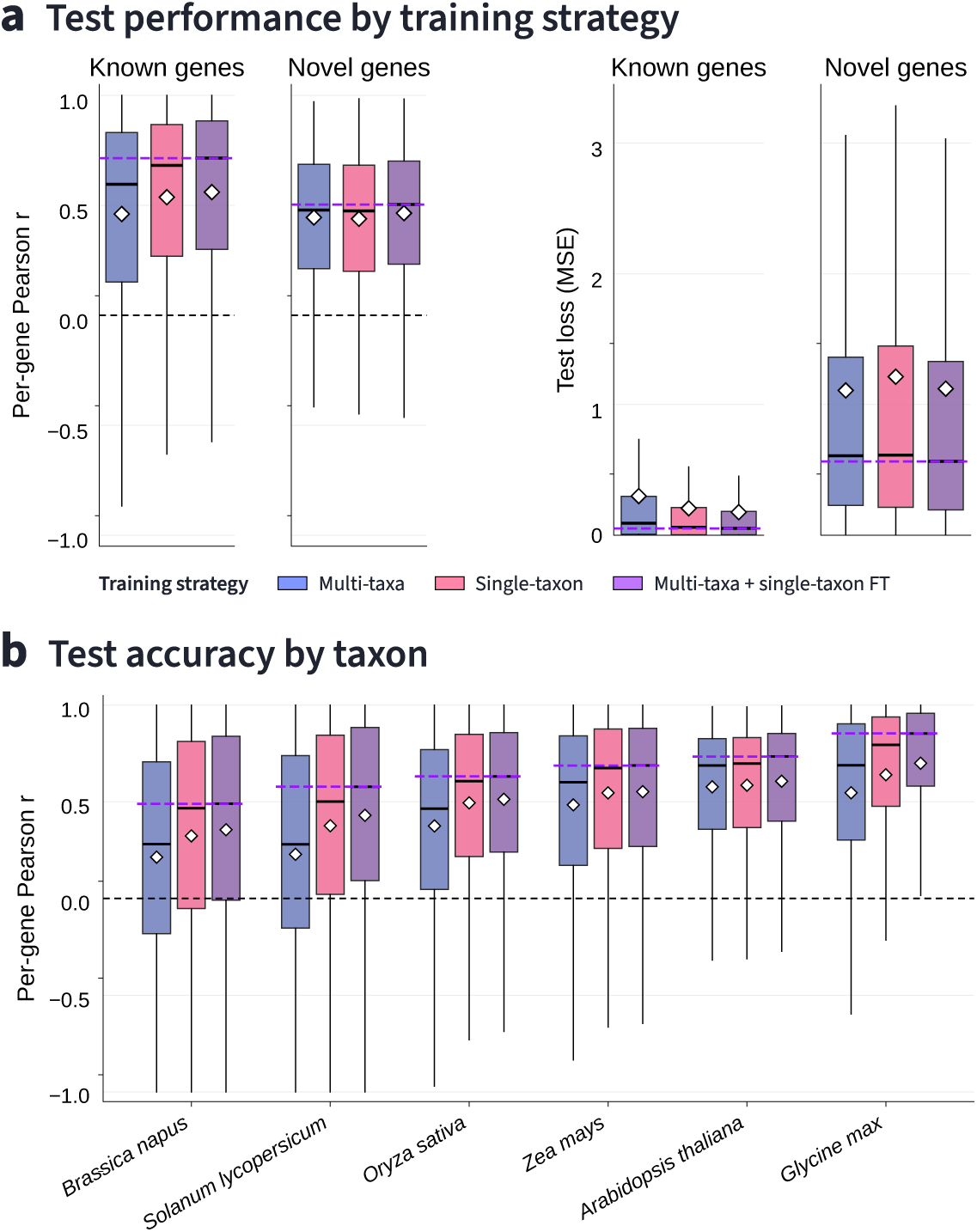
Multi-species pretraining improves taxon-specific prediction after fine-tuning. **a**, Test-set distributions of per-gene Pearson correlation and MSE for a shared six-taxon model (blue), taxon-specific models trained from scratch (pink), and the shared model subsequently fine-tuned on each taxon (purple). Known and novel genes are shown separately. Purple dashed lines mark the fine-tuned model’s median; white diamonds denote arithmetic means. **b**, Per-gene correlation distributions for each taxon, pooling the known- and novel-gene groups. FT denotes fine-tuning. Boxplot conventions follow Fig. 3. All strategies were trained to convergence, and all trainable parameters were updated during

The benefit extended to genes absent from training. For novel genes, fine-tuning achieved mean correlation 0.465, compared with 0.439 for taxon-specific training from scratch and 0.445 for the shared model, alongside the lowest median MSE. Per-taxon distributions show how these aggregate results vary across the six species (Fig. 5b).

Learning across species therefore provides a useful starting point for crop-specific expression prediction. Subsequent fine-tuning improves on both the shared model before fine-tuning and training from scratch on the target taxon, supporting a training strategy that combines broad regulatory evidence with taxon-specific refinement.

### Sparse edits produce strong, context-selective predicted responses

Given CROP-1’s ability to predict context-dependent expression, we asked whether it could guide cis-regulatory edits that alter a gene’s activity selectively in a chosen tissue or condition. This is a central requirement for crop engineering: an expression change should occur where it supports the intended trait while preserving activity in other contexts to reduce pleiotropic effects. We tested this capability by targeting either up- or down-regulation in one context and maintenance of baseline expression in four others.

A Ledidi-based optimiser^37^ adjusted the DNA input while keeping CROP-1 and the TF contexts fixed (Fig. 6). Backpropagating the expression objective through CROP-1 guides sequence updates across editable positions simultaneously, providing a more directed search than random mutation. It can also reduce the repeated candidate evaluations needed by prediction-only evolutionary approaches^37^. Substitutions were restricted to the 500 nucleotides immediately upstream of the annotated ATG (i.e., the translation initiation site) within the 5-kb prediction window. For most genesin these panels, thisinterval coversboth promoter sequence and the 5′ untranslated region. The optimiser balances the expression objective with a sequence-change penalty and can adjust that penalty to search within a desired substitution budget. We examined candidates across penalty settings, grouping them by their realised number of substitutions.

**Figure 6.**
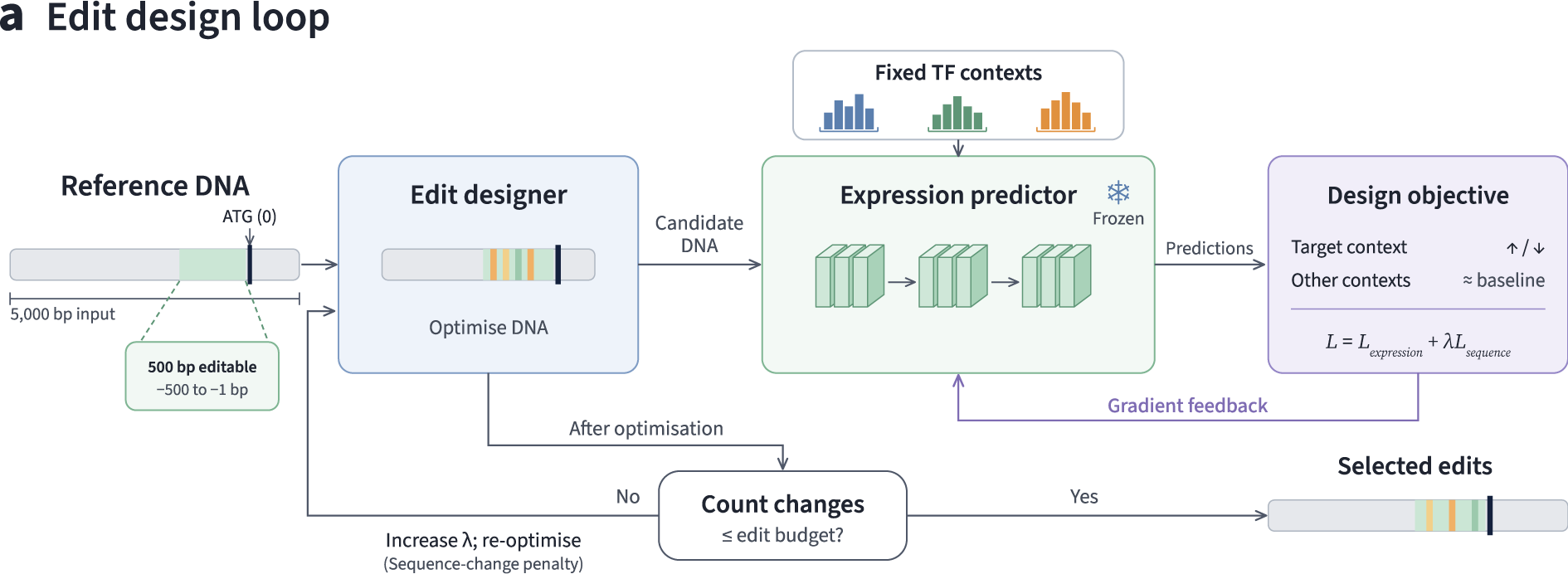
A differentiable workflow for context-selective sequence editing. **a**, A Ledidi-based designer optimises the 500-bp interval immediately upstream of the annotated ATG (−500 to −1) within the 5,000-bp input sequence, while the trained CROP-1 predictor and supplied TF contexts remain fixed. The objective combines a desired increase or decrease in the target context, preservation of baseline expression in the other contexts and a sequence-change penalty weighted by *λ*. Gradients guide sequence optimisation; the resulting substitutions are counted, and the penalty can be increased before re-optimisation to meet the desired edit budget. The schematic shows three TF contexts for clarity; the soybean and tomato design experiments each use five. Predicted outcomes are reported in Fig. 7 and Supplementary Fig. 1, respectively.

Small edit sets produced strong, context-selective predicted responses in soybean (Fig. 7a,b). Across ten genes and five contexts—dry seed, endosperm, epicotyl, flower bud and root—designs with 1–10 substitutions achieved mean target responses of +1.70 log2 fold change for up-regulation and −2.34 for down-regulation. With 11–20 substitutions, the responses reached +2.83 and −4.12. The corresponding non-target responses were much smaller, at +0.31 and −0.36, measured as the mean signed median across the four other contexts. The full context matrices show the distribution of these effects across tissues.

**Figure 7.**
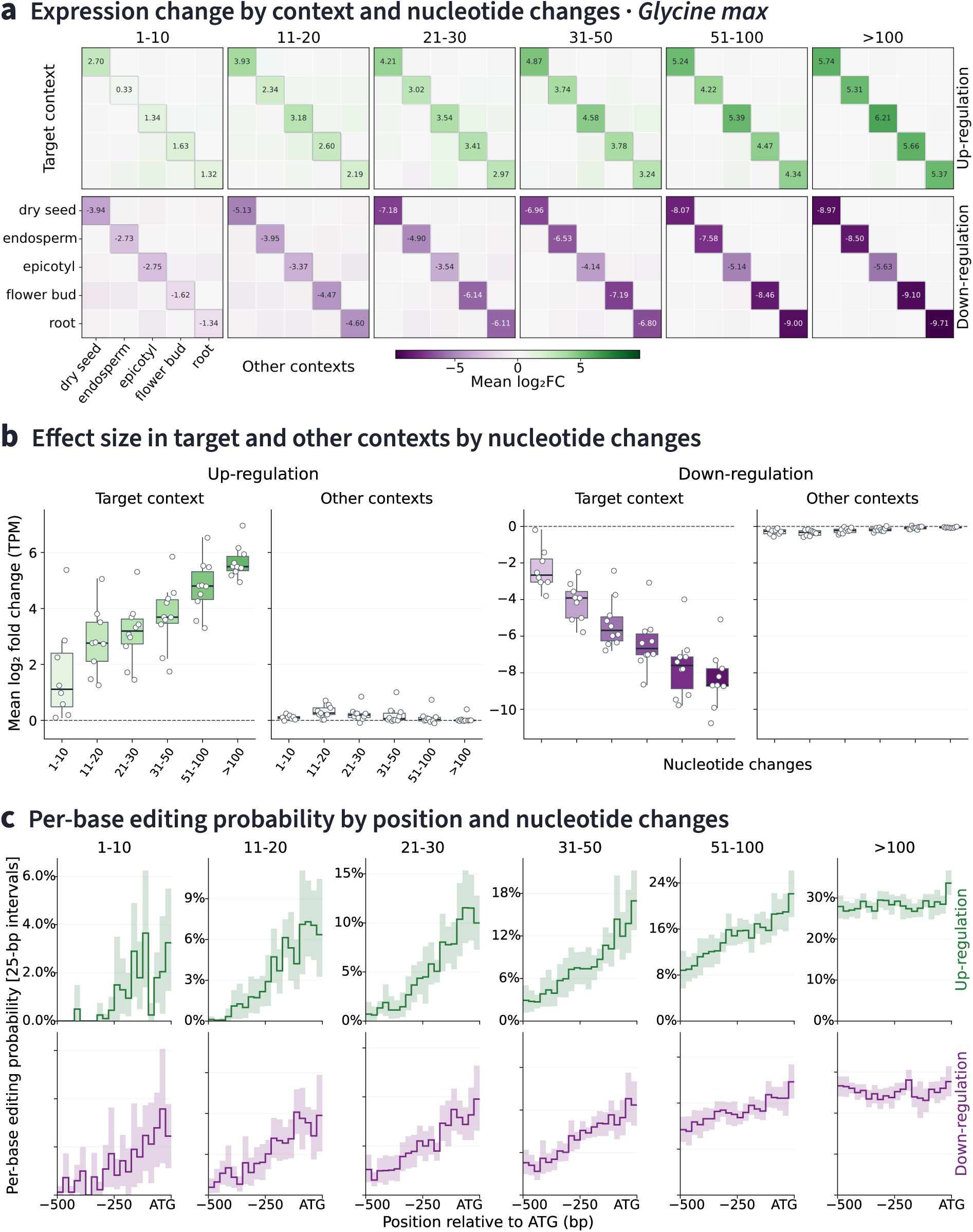
Sparse sequence edits produce context-selective predicted expression changes in soybean. Designs were generated for ten genes across dry seed, endosperm, epicotyl, flower bud and root using the workflow in Fig. 6. **a**, Mean predicted log2 fold changes, grouped by realised substitution count (1–10, 11–20, 21–30, 31–50, 51–100 and more than 100) and requested direction of regulation. Rows denote target contexts and columns denote the contexts in which expression is evaluated; labelled diagonal entries show target-context responses and off-diagonal entries show responses in the other contexts. **b**, Gene-level distributions of target responses and signed median responses across the four non-target contexts. Each point represents one gene after averaging across its available target contexts within a substitution-count bin. Boxes show the interquartile range, centre lines the median and whiskers extend to 1.5 interquartile ranges; dashed lines mark zero change. **c**, Mean per-base editing frequency in 25-bp intervals from −500 to −1 relative to the A of ATG, shown separately for up-regulation (green) and down-regulation (purple). Shading shows pointwise 95% percentile intervals from 5,000 whole-gene bootstrap resamples. Expression predictions were converted to TPM before calculating log2 fold changes; effect summaries use equal-gene weighting (Methods).

Tomato designs showed the same broad pattern of selective predicted expression changes (Supplementary Fig. 1a,b). Across eight genes evaluated in fruit, leaf, pollen tube, root and stem, designs with 11–20 substitutions achieved mean target responses of +4.15 and −3.57 log2 fold change, with corresponding non-target responses of +0.23 and −0.28 (Table 3). In both crops, larger edit sets generally produced stronger target responses, while average changes outside the target context remained comparatively small.

**Table 3.** Predicted effects of sparse designs in soybean and tomato. Only designs with 1–20 substitutions are summarised here. Values are mean target log2 fold changes and means of the per-design signed median across four non-target contexts, using equal-gene weighting. Up and down refer to the requested direction in the target context.

| Species | Substitutions | Up |  | Down |  |
| --- | --- | --- | --- | --- | --- |
|  |  | Target | Other contexts | Target | Other contexts |
| Soybean | 1–10 | +1.70 | +0.11 | –2.34 | –0.29 |
|  | 11–20 | +2.83 | +0.31 | –4.12 | –0.36 |
| Tomato | 1–10 | +1.45 | +0.28 | –2.11 | –0.23 |
|  | 11–20 | +4.15 | +0.23 | –3.57 | –0.28 |

Sparse substitutions were preferentially located near the translation start in both crops (Fig. 7c; Supplementary Fig. 1c). For soybean designs with 1–10 substitutions, mean per-base editing frequency in the ATG-proximal 25-bp interval (−25 to −1) was 3.25% for up-regulation and 2.45% for down-regulation, compared with 0% and 0.125% in the distal interval (−500 to −476). Tomato showed the same proximal enrichment, and larger edit sets in both crops were distributed more broadly. This pattern may reflect the location of regulatory elements in the promoter and 5′ untranslated region, together with the model’s sensitivity to those sequences. The prioritised sites provide concrete candidates for testing whether changes near the translation start can produce the predicted target-context response while preserving expression elsewhere. Measuring these effects across the queried contexts, alongside edits at more distal positions, would establish whether the predicted enrichment corresponds to biological sensitivity in this region.

Across soybean and tomato, CROP-1-guided sequence optimisation identified small substitution sets that strongly increased or decreased predicted expression in the specified target context, with much smaller average responses in the non-target contexts. These results show that a shared sequence-and-context predictor can guide sparse edits towards both the requested target response and preservation of expression elsewhere, providing concrete candidates for experimental tests of context-selective regulation in endogenous crop genes.

## Discussion

CROP-1 makes biological state an explicit input to plant genomic modelling, so a shared architecture can represent the same cis-regulatory sequence across contexts and guide edits towards a desired expression pattern. This capability addresses a practical crop-engineering challenge: changing gene activity where it supports a trait while limiting changes in other tissues, developmental stages or conditions, thus minimising undesired pleiotropic effects. Across six plant taxa, TF context improved prediction beyond sequence-only comparators, with particularly strong gains in gene-level prediction in novel contexts. Larger models and broader experiment coverage delivered further improvements, while tissue aggregation and multi-species pretraining followed by taxon-specific fine-tuning improved the use of available data. Paired with gradient-based sequence optimisation, CROP-1 guided sparse edits in soybean and tomato with strong, context-selective predicted effects, connecting advances in expression prediction to the design of endogenous regulatory alleles.

### TF context improves prediction across biological states

Explicit TF context enables CROP-1 to predict how plant gene expression varies across biological states and to propose sequence edits with selective predicted effects. The strongest evidence comes from two distinct comparisons: at approximately 100M parameters, adding TF context improved prediction over the sequence-only CROP-1 counterpart; the same 100M context-aware model also achieved higher mean correlations than the approximately 320M Botanic1-S model after full fine-tuning on the same expression task. The gains in gene-level correlation were especially pronounced in novel taxon–tissue contexts, for both known and novel genes (Fig. 3b). Increasing sequence-model capacity alone therefore does not account for the gains observed here. Explicit TF inputs contribute information about the state in which a sequence is active, and allow a shared predictor to respond to changes in that state.

These gains are most apparent when evaluation separates context-associated variation from baseline differences between genes. Tissue-level correlations before gene-median subtraction were high across all tested models. Centring exposed differences in their ability to predict expression relative to each gene’s typical level, consistent with the advantage of CROP-1 in the complementary within-gene analysis. For applications that depend on selective expression changes, these metricsare more informative than cross-gene correlation alone. The results support evaluating both axes whenever a model is intended to guide context-specific interventions.

The TF representation provides a biological basis for this behaviour. Motifs describe potential DNA recognition, while TF expression supplies a measurable indicator of regulatory state. Binding also depends on interactions among regulators and on chromatin: cooperative motif arrangements can influence occupancy^38^, TF pairs can recognise composite sites with altered sequence preferences^39^, and pioneer factors can recognise partial motifs exposed on nucleosomes^40^. In plants, promoter features contribute differently across assay systems^15^ and enhancer elements can interact cooperatively in the light but more additively in the dark^16^. These findings motivate treating regulatory activity as a joint property of sequence and biological state. Understanding which aspects of the TF context drive CROP-1’s predictions, and how their influence depends on the input sequence, will require further investigation through feature attribution and targeted *in silico* perturbation analyses.

The choice of context input determines which aspects of regulatory state a model can use. GET combines sequence-derived features with chromatin accessibility^17^, whereas Corgi and RegFM use regulator expression or activity to condition sequence representations^18,19^. These human models provide precedents for supplying biological state explicitly, but they draw on different measurements of that state. CROP-1 uses TF motifs and expression, allowing it to draw on the broad coverage of plant transcriptomic resources without requiring matched chromatin-accessibility assays. Adding accessibility or protein-level measurements could capture regulation that TF RNA abundance alone does not resolve; evaluating such additions would establish whether their predictive benefit justifies the extra data requirements for a given crop and condition.

Explicit context and genomic pretraining should also be complementary. PlantCAD2 and BOTANIC-1 learn broadly useful plant sequence representations^11,13^, while Nucleotide Transformer v3 connects sequence learning to functional supervision across species^12^. Combining such representations with TF-context inputs could bring broader sequence knowledge to the prediction of state-dependent responses. CROP-1’s performance on novel taxon–tissue combinations motivates testing this approach in additional developmental and environmental conditions. Because all conditions use the same TF-context encoding framework and scalar-output predictor, CROP-1 can accommodate an arbitrary number of queried conditions without adding condition-specific heads or increasing the number of model parameters. Measured TF expression profiles can therefore extend the set of queried states, with evaluation in those states establishing how well the learned sequence–context relationships transfer.

### Scaling and fine-tuning improve gene expression prediction in plants

The capacity and data scaling experiments show that context-aware prediction can improve systematically as training resources expand. Increasing parameters from 4M to 400M reduced held-out set error (validation MSE) by 47.8%, and expanding experiment-level supervision reduced it by 57.7%. Both series continued to improve at their largest tested scales. Their empirical power laws connect plant expression modelling to established observations in neural scaling^22,23^. Bahri et al. distinguish scaling driven by reduced variability from finite data or model width from scaling driven by finer resolution of an underlying data manifold—the structure occupied by the inputs^41^. In the latter regime, more data cover that structure more densely, while greater capacity allows the model to represent finer distinctions. For CROP-1, this suggests a hypothesis in which broader sampling of TF expression profiles and greater capacity resolve increasingly detailed sequence–context relationships. Our fits do not identify a particular theoretical regime, but they provide quantitative evidence for further gains from capacity and expression coverage.

Data composition determines how efficiently that capacity can be trained. Tissue aggregation retained substantial predictive accuracy on individual experiments with roughly 40-fold fewer training observations. This suggests that repeated measurements within a tissue contain enough shared signal to support compact supervision, although the present comparison does not separate noise reduction from coverage of common regulatory states. The additional gains from experiment-level training show the value of retaining finer contextual variation. Together with evidence for informative data selection in machine learning^24^, these findings motivate starting with broad tissue coverage and then adding experiments that contribute distinct TF expression profiles or responses. The clustering analysis provides one way to organise that expansion.

Multi-species learning improves the use of existing measurements in a different way: it supplies a starting representation that can be refined for an individual crop. Taxon-specific fine-tuning outperformed both the shared model and converged models trained from scratch on the target taxon, supporting transfer within the six-species training collection. This result complements evolutionary approaches to plant expression prediction^20^ and cross-species genomic pretraining^21^. A natural extension is to fine-tune the shared model for a defined breeding use case, such as a developmental transition or stress response, using measurements from the relevant conditions. Testing taxa with limited data, holding out entire taxa and separating related gene families would establish how far that transfer extends beyond the settings evaluated here. By representing species-specific TF repertoires through a common motif–expression input format, CROP-1 can be trained jointly across species without changing the model architecture or adding species-specific prediction heads.

### Context-specific edits for crop trait engineering

Context-selective edit design is valuable because agronomic genes often affect more than one trait^3^. Regulatory editing has already generated quantitative trait variation^1^, improved maize performance under drought^2^ and separated competing effects on rice yield components^3^. Lou et al. extended this principle to environmental responsiveness: inserting heat-shock elements into cell-wall-invertase regulatory regions improved source–sink allocation and heat-stress resilience in tomato and rice^4^. This demonstrates the agronomic value of coupling expression to environmental conditions. Extending these successes to new genes and conditions requires a way to choose among many possible regulatory edits and anticipate their effects beyond the intended context. A model that jointly considers target and non-target responses can prioritise alleles with the desired expression pattern before construction and testing. Paired with a gradient-based sequence optimiser, CROP-1 guided the design of cis-regulatory edits with 11–20 substitutions in soybean and tomato that produced strong predicted target responses, while the average response across the other queried contexts remained much smaller. The same design procedure could generate candidates for both up- and down-regulation.

Our approach to edit design builds on experimental advances in model-guided regulatory sequence engineering. Random promoter libraries in yeast have revealed learnable sequence–activity relationships^25^ and supported the engineering of promoter strength^27^; cell-type-selective elements in mammals and tissue-specific enhancers in Drosophila demonstrate that biological specificity can also be a design objective^28,29^. Differentiable methods such as Ledidi make it possible to optimise sequence inputs directly against these objectives^37^. Here, that search was applied to endogenous plant cis-regulatory sequences under substitution constraints. The resulting edit candidates address a concrete crop-engineering question: which small changes could alter expression in a desired context while limiting changes elsewhere?

The next step is to measure those edit candidates in the intended target and non-target contexts. Their reported effects are predictions from the model used for optimisation, so prospective testing is needed to establish biological selectivity. Plant promoter assays provide a precedent for model-guided design and measurement^15^. Jevtic et al. combined a multi-species genomic model with a protoplast-based massively parallel reporter assay to identify tomato promoter variants with increased activity^42^. For CROP-1-designed candidates, reporter assays across the queried contexts could establish which edits warrant testing at endogenous loci.

Experimental panels could include different substitution patterns with similar predicted effects, helping distinguish alternative sequence routes to the same expression objective. Adaptive sampling offers a way to favour promising sequences in regions where predictions are better supported by training data^43^. Methods developed for model-guided selection could also provide uncertainty intervals that account for how the predictor influenced candidate choice^44^. Feeding measurements from these tests back into training would improve subsequent design rounds.

Small substitution sets also offer a practical connection to precise genome editing. Combining a knockout with tissue-specific complementation can restrict gene activity, as demonstrated by vessel-specific restoration of *CCR1* expression in Arabidopsis, which reduced lignin while preserving biomass yield^45^. That strategy requires a suitable promoter and an introduced expression cassette. Sparse regulatory editing, on the other hand, modifies the endogenous regulatory sequence while retaining the native coding region and most of its surrounding regulatory architecture. Prime editing supports specified sequence changes^30^, with demonstrations in rice and wheat^31^, maize^46^ and tomato^47^ establishing its applicability across crop genomes. Incorporating editing-system constraints into optimisation could help prioritise candidates that combine a useful predicted expression profile with a feasible construction strategy. Explicit substitution budgets also connect model-guided design to the European framework for new genomic techniques. Designs intended to meet NGT category 1 criteria, for example, require assessment of the resulting plant against all applicable requirements–notably limits on the number and size of genetic modifications^32,33^.

By making target and non-target context part of the same expression optimisation objective, CROP-1 provides a way to search for endogenous cis-regulatory alleles with effects concentrated in the tissues, developmental stages and conditions where they are needed. Beyond edit design, modelling sequence–state relationships could also help identify and prioritise genes whose modulation affects agronomic traits in particular contexts and cultivars. Together, target prioritisation and context-selective edit design could support improvements in disease resistance or stress tolerance while limiting adverse effects on yield, flavour and developmental timing.

## Methods

### Expression resource and benchmark definition

Expression data were obtained as pre-quantified transcripts-per-million (TPM) tables from PlantExp^34^ for *Arabidopsis thaliana*, *Brassica napus*, *Glycine max*, *Oryza sativa* Japonica Group, *Solanum lycopersicum* and *Zea mays*. Experiments were selected using curated cultivar and genotype metadata, and expression gene identifiers were matched to the corresponding reference annotation. RNA-seq alignment and quantification were performed by the source resource; our processing began with its quantified expression tables.

The corresponding reference assemblies were TAIR10, AST_PRJEB5043_v1, Glycine_max_v2.1, IRGSP-1.0, SL3.0 and B73_RefGen_v4, respectively, with gene annotations obtained through Ensembl Plants^48^. Prediction inputs were 5,000-bp genomic windows comprising 3,000 bp upstream of the annotated gene boundary and 2,000 bp extending downstream from and including that boundary. The anchor depended on gene orientation: the annotated start coordinate for genes on the forward strand and the annotated end coordinate for genes on the reverse strand. This annotated 5′ gene boundary served as the TSS reference. Upstream and downstream were defined relative to the direction of transcription, and sequences were oriented accordingly. No representative transcript was selected. Windows extending beyond reference-sequence boundaries were padded with ambiguous nucleotide bases. These prediction windows were defined separately from the translation-start-relative intervals used for computational edit design.

Tissue labels were harmonised across experiments. For each gene–taxon–tissue combination, we transformed the experiment-level TPM values using the natural logarithm, log1p(TPM), and took the median across experiments to obtain the tissue-level expression target. Missing measurements were excluded and measured zeros retained. Each experiment contributed directly to this median, without prior aggregation into biological or technical replicate groups or equal weighting of studies.

Tissue-level aggregation preceded partitioning. The test set combined held-out taxon–tissue combinations, gene-level holdouts and additional held-out observations; training comprised the remaining gene–taxon–tissue combinations. Gene novelty was defined by gene-identifier exposure in training, and context novelty by exposure to taxon–tissue combinations in training target supervision. All six taxa and all 234 tissue descriptors were represented in training, with selected taxon–tissue combinations held out.

The tissue-aggregated tables contain 14,459,659 training observations and 910,725 test observations, with no shared gene–taxon–tissue triples. There are 308,898 distinct training genes and 195,171 test genes, including 5,942 novel genes. Their union contains 314,840 genes. The resource spans 234 tissue descriptors and 357 taxon–tissue combinations. The experiment-level dataset and its construction are described below under ‘Experiment selection and taxon-specific fine-tuning’.

### TF context and model architecture

CROP-1 is a convolutional genomic model that predicts one gene-expression value from a DNA sequence and its TF context. A one-hot encoded nucleotide representation enters a convolutional backbone with residual blocks that extract sequence features at successive depths. Pooling condenses these features before a feed-forward head produces the scalar expression prediction.

TF identities and associated DNA-binding motifs were obtained from PlantTFDB^35,36^ and linked to reference gene identifiers and expression measurements. The resource includes experimentally supported and computationally transferred motif assignments. Binding motifs were represented as nucleotide-probability matrices and padded to a common length, using uniform nucleotide probabilities for padding. Each TF retained the same motif representation across contexts, while its expression varied by experiment or tissue. Tissue-level TF expression profiles used the same median-of-log-transformed-TPM procedure as the target labels.

Each TF is represented by a token combining its fixed-length motif representation with its measured expression in the requested context. The full set of these motif–expression tokens constitutes the TF context. The number of TF tokens can vary across taxa, and masks identify unavailable inputs. When the prediction target is itself a TF, its expression value is withheld and its token is excluded from attention, preventing direct leakage of the target expression value through its own TF-context entry. Evaluation with TF context assumes access to measured TF expression in the queried context.

After removal of all-zero TF expression profiles, the experiment-level resource contains 10,704 eligible TF expression profiles for Arabidopsis, 243 for Brassica, 1,073 for soybean, 1,679 for rice, 146 for tomato and 3,089 for maize. The tissue-level inventories of TF genes with annotated binding motifs in PlantTFDB contain 620, 617, 370, 269, 278 and 250 TF genes, respectively.

Cross-attention integrates TF context with the sequence representation at multiple depths. Sequence features supply attention queries, and TF tokens supply keys and values, allowing local sequence features to draw on the relevant regulatory context. The attention output is combined with the sequence features through residual connections and feed-forward processing. The same TF context is reused across depths, with separate learned projections and attention parameters in each block. TF tokens do not undergo self-attention with one another. We varied model capacity within this architecture family; Table 4 reports the parameter counts used for the scaling analysis.

**Table 4.** Parameter counts of the CROP-1 scaling models. Rounded labels identify models throughout the text and figures; total parameter counts include embeddings.

| Model label | Total parameters |
| --- | --- |
| 4M | 4,000,004 |
| 10M | 10,046,420 |
| 40M | 38,435,512 |
| 100M | 101,411,782 |
| 400M | 400,956,330 |

### Training and scaling analysis

The model and training code were implemented in PyTorch and Lightning. CROP-1 training used distributed computation on NVIDIA GH200 GPUs in the Isambard-AI Phase 2 cluster^49^. Models were optimised with AdamW using learning-rate warm-up followed by validation-based learning-rate reductions, mixed bfloat16 precision and gradient clipping. The parameter-scaling, experiment-data and taxon-specific fine-tuning models were trained to convergence of validation loss, and the checkpoint with the lowest validation MSE was selected for evaluation. The parameter-scaling series used a common optimisation schedule on four nodes with four GPUs per node (16 GPUs in total), with 128 observations per GPU and a global batch size of 2,048. Across model sizes, the selected checkpoints occurred at 185,000–322,500 optimiser updates, equivalent to approximately 28–48 epochs over the training data.

For the tissue-aggregated parameter-scaling series, a fixed validation partition of 722,982 observations (5% of the training partition) was reserved for model selection, leaving 13,736,677 observations for fitting. Models minimised mean squared error over observed expression targets. Test observations were used for the reported generalisation analyses.

We used the single-variable power-law form of Kaplan et al.^22^, *L*(*X*) = (*X_c_*/*X*)^*α*^, to describe validation MSE as a function of model or dataset size. For our analysis, we fitted this form by least squares in log space to the selected checkpoint losses from the five converged model-size runs or five converged dataset-size runs. Here, *X* denotes total parameter count or the realised number of training observations, *X_c_* is a fitted scale constant and *α* is the scaling exponent. Fit quality was assessed by R^2^ in log space.

Training compute was estimated from an initial profile of each run using PyTorch’s operation counter. Profiling covered the training operations, including forward and backward passes, through the first optimiser update. The counted operations and processed examples were summed across distributed workers to estimate FLOPs per training example. Cumulative FLOPs were then calculated by multiplying this estimate by the cumulative number of examples processed across workers, with counters retained when training resumed. These estimates cover operations supported by the counter and were aligned with the validation-loss history to produce the compute-scaling plots.

### Sequence-only and pretrained comparators

The CROP-1 comparator without TF context has approximately 100 million parameters, matching the scale of the context-aware model. It retains the convolutional sequence backbone, removes TF cross-attention and adjusts backbone capacity to preserve a comparable total parameter count. A multitask head predicts expression for 234 tissue descriptors. Training examples correspond to genes with their available tissue-expression labels, and missing labels are masked in the MSE objective. A validation subset is used for checkpoint selection.

We evaluated Botanic1-S, NTv3-pre and PlantCAD2-Small, selecting available model-size variants as close as possible to the parameter count of the 100-million-parameter CROP-1 model. These variants contain approximately 320 million, 108 million and 90 million parameters, respectively (Table 1). Each pretrained backbone was fully fine-tuned on the same expression task with a feed-forward regression head producing the tissue-expression outputs.

All comparators received the same 5-kb genomic windows with the same transcriptional orientation described above. For each pretrained model, the input was encoded using its native sequence representation, and the final backbone features were mean pooled across sequence positions to form an input to the tissue-expression regression head. Botanic1-S used its final normalised sequence features, excluding the classification token from pooling. PlantCAD2 produced paired forward and reverse-complement feature representations; the reverse-complement branch was realigned, and each branch was pooled separately and passed through the same regression head before averaging the two predictions. NTv3 used its final nucleotide-resolution features after deconvolution; sequences were right-padded from 5,000 to 5,120 positions to satisfy the backbone’s input requirements, with the 120 padding positions excluded from pooling. The regression heads produced one value for each of the 234 tissue descriptors, and missing expression labels were masked during training. No additional reverse-complement augmentation was applied.

All three pretrained comparators used the same fine-tuning hyperparameters and validation-based learning-rate schedule. Both the backbone and regression head were updated with AdamW at an initial learning rate of 5 × 10^−5^, weight decay of 0.01 and an effective batch size of 2,048 genes, using gradient accumulation where required. Training used mixed bfloat16 computation, float32 loss reductions and gradient-norm clipping at 0.5. The learning rate was reduced by a factor of 0.25 after validation loss plateaued, with scheduler patience set to two validation checks and a minimum learning rate of 10^−6^. The 2,780-gene validation set was evaluated in full every 250 optimiser updates and at epoch end. Training continued until validation improvement plateaued; the lowest-validation-MSE checkpoints used for evaluation were reached after approximately 7–14 epochs. All models were evaluated on the same gene/context prediction task using the metric definitions below.

### Experiment selection and taxon-specific fine-tuning

Experiment-level observations were aligned to the train/test partition defined at tissue level using gene–taxon–tissue keys, yielding 568,566,061 training rows and 48,179,058 test rows with no shared keys. A fixed validation sample of 262,144 observations was shared across the experiment-data scaling runs; removing these observations left 568,303,917 observations in the complete experiment-training dataset. Different target genes from the same experiment can occur in different partitions; the test is defined by held-out gene–taxon–tissue combinations.

Within each taxon, experiments were represented by their TF expression profiles. Missing or zero-variance features were removed before clustering, leaving expression values for 620, 615, 369, 269, 276 and 250 TFs in Arabidopsis, Brassica, soybean, rice, tomato and maize, respectively. These counts refer to the TFs retained for clustering, with one expression feature per TF; they are distinct from the feature dimension of an individual motif–expression token used by CROP-1.

Complete-linkage hierarchical clustering used one minus Pearson correlation as the distance between experiment profiles. We explored a broader range of distance thresholds and selected 0.5, 0.25, 0.15 and 0.025 to obtain subsets approximating the desired 1%, 5%, 10% and 50% context fractions, respectively. The full eligible dataset provided the 100% condition. Each cluster was represented by the experiment with the smallest sum of distances to the other members. Ties were resolved deterministically. If that representative was not training eligible, an eligible member was selected; clusters without eligible members were excluded. All eligible expression observations for each selected representative were retained apart from the fixed validation sample.

From an inventory of 16,934 experiment-level TF expression profiles across the six taxa, the four thresholds initially selected 215, 852, 1,825 and 8,891 representatives and yielded 210, 810, 1,693 and 8,303 training contexts after eligibility handling. The training-eligible pool contained 15,899 experiment contexts; clustering used the full profile inventory before this eligibility restriction was applied, and subsets need not be nested. For visualisation, t-SNE was fitted independently within taxa to standardised TF expression features using seed 42 and perplexity 30.

Taxon-specific fine-tuning experiments used the 100M model and compared a single shared model trained jointly on the complete experiment data from all six taxa, taxon-specific models trained from scratch using only data from the corresponding taxon, and taxon-specific fine-tuning from the shared checkpoint. All trainable parameters were updated during taxon-specific fine-tuning, including those in the sequence backbone, TF-context modules and prediction head. Within each taxon, scratch training and fine-tuning used the same target-taxon training data. The strategies used the same training hyperparameters and were trained to convergence of validation loss, and the lowest-validation-MSE checkpoint was selected. Training to convergence avoided imposing a common update limit across differently sized datasets; the comparison did not match total pretraining and fine-tuning compute.

### Evaluation and statistical analysis

Tissue-level metrics compare predictions and targets across genes within each tissue descriptor, which can include multiple taxa. Gene-level metrics compare predictions and targets across available contexts for each gene. Within each generalisation stratum, Pearson correlation and MSE were calculated for the evaluated genes or tissue descriptors and combined as unweighted means.

For the external-model comparison in Fig. 3, the same explicit training-gene and novel-gene lists defined the groups for all six models. For the non-stratified results, all test observations were analysed without separating them by gene or context novelty. We calculated one correlation across contexts for each gene and one across genes for each tissue descriptor, then summarised these distributions by their means and medians, weighting each gene or tissue descriptor equally.

Except for the explicitly uncentred tissue-level comparison in Fig. 3a, tissue-level Pearson correlations were calculated after subtracting a gene-specific median from both predictions and observations. This median was calculated across the available tissue-aggregated training and test labels for that gene and used only as an evaluation reference. This gene-median centring removes the baseline expression differences between genes: without it, a predictor that assigns each gene its average expression across contexts can achieve a high cross-gene Pearson correlation while failing to capture context-specific responses. Correlation after centring therefore evaluates how well predictions capture expression relative to each gene’s characteristic level. Subtracting the same reference from predictions and observations leaves individual squared errors, and hence MSE, unchanged. It also leaves within-gene correlation unchanged because the reference is constant across contexts for a given gene.

Experiment-level evaluation computes per-gene Pearson correlation and MSE across experiment contexts using predictions and observed expression on the log1p(TPM) scale, without tissue-level aggregation or subtraction of a tissue-specific expression baseline. Known and novel groups follow each model’s actual training exposure. Boxplots show distributions across genes or tissue descriptors: central lines denote medians, boxes span the interquartile range, whiskers extend to 1.5 interquartile ranges and diamonds denote arithmetic means. Outlier markers are suppressed for readability, without removing those values from the mean.

### Context-specific sequence editing

We used a gradient-based sequence designer based on Ledidi^37^ to search for edits with context-selective predicted effects. The trained CROP-1 predictor and the TF contexts were held fixed while the editable DNA sequence was optimised. For each gene, predictions for the unmodified sequence defined the baseline expression in each context. The design objective combined a requested increase or decrease in one target context with preservation of expression near baseline in the remaining contexts. A sequence-change penalty balanced the requested expression response against the number of substitutions. Adjusting this penalty allowed the sequence optimiser to search for edit candidates within a specified nucleotide substitution budget; the achieved number of nucleotide changes was counted for each edit candidate.

Positions −500 to −1 relative to the A of the annotated ATG were editable, with the rest of the 5-kb sequence held fixed. For most genes in these panels, the editable interval includes both promoter sequence and the 5′ untranslated region. Candidates generated across sequence-penalty settings were grouped by their realised substitution count: 1–10, 11–20, 21–30, 31–50, 51–100 and more than 100. The soybean panel contained ten genes across dry seed, endosperm, epicotyl, flower bud and root; the tomato panel contained eight genes across fruit, leaf, pollen tube, root and stem. Each gene was targeted for both up- and down-regulation in every context.

### Design effects and positional analysis

CROP-1 expression outputs, which are on the log1p(TPM) scale, were converted back to TPM before calculating log2 fold changes between edited and unmodified sequences. Effects were first averaged across designs for the same gene, target context and substitution-count bin, then across available target contexts within each gene. The reported cross-gene means gave equal weight to each gene. For non-target effects, each design was first represented by the signed median response across its four non-target contexts. Heatmaps retained the full target-by-context comparison, again using equal-gene weighting.

For position analysis, changed nucleotides were mapped to the ATG-relative coordinate system. Counts in each 25-bp interval were divided by the interval length to obtain per-base editing frequency, followed by the same hierarchical averaging described above. Uncertainty was estimated using 5,000 whole-gene bootstrap resamples. Each resample drew genes with replacement, retaining the original number of genes in the panel and keeping all contexts, candidates and positions associated with each selected gene together. We recomputed the mean editing frequency for each interval in every resample, preserving equal-gene weighting. The 2.5th and 97.5th percentiles of the resulting bootstrap distribution defined the pointwise 95% interval at each position bin.

## Data availability

The expression data used in this study are publicly available from PlantExp^34^. Reference genome assemblies and gene annotations are available from Ensembl Plants^48^. TF identities and binding motifs were obtained from PlantTFDB^35,36^. The species, assembly versions and preprocessing procedures are described in Methods.

## Author contributions

**Conceptualisation, project administration, supervision, and funding acquisition:** P.A., D.H.

**Data curation:** D.S., J.D.M., P.A.

**Investigation, software and methodology:** P.A., L.W., J.D.M.

**Formal analysis and writing:** P.A.

## Acknowledgements

We thank members of Biographica Ltd. for their review and comments on the manuscript.

## Supplementary Figures

**Supplementary Figure 1.**
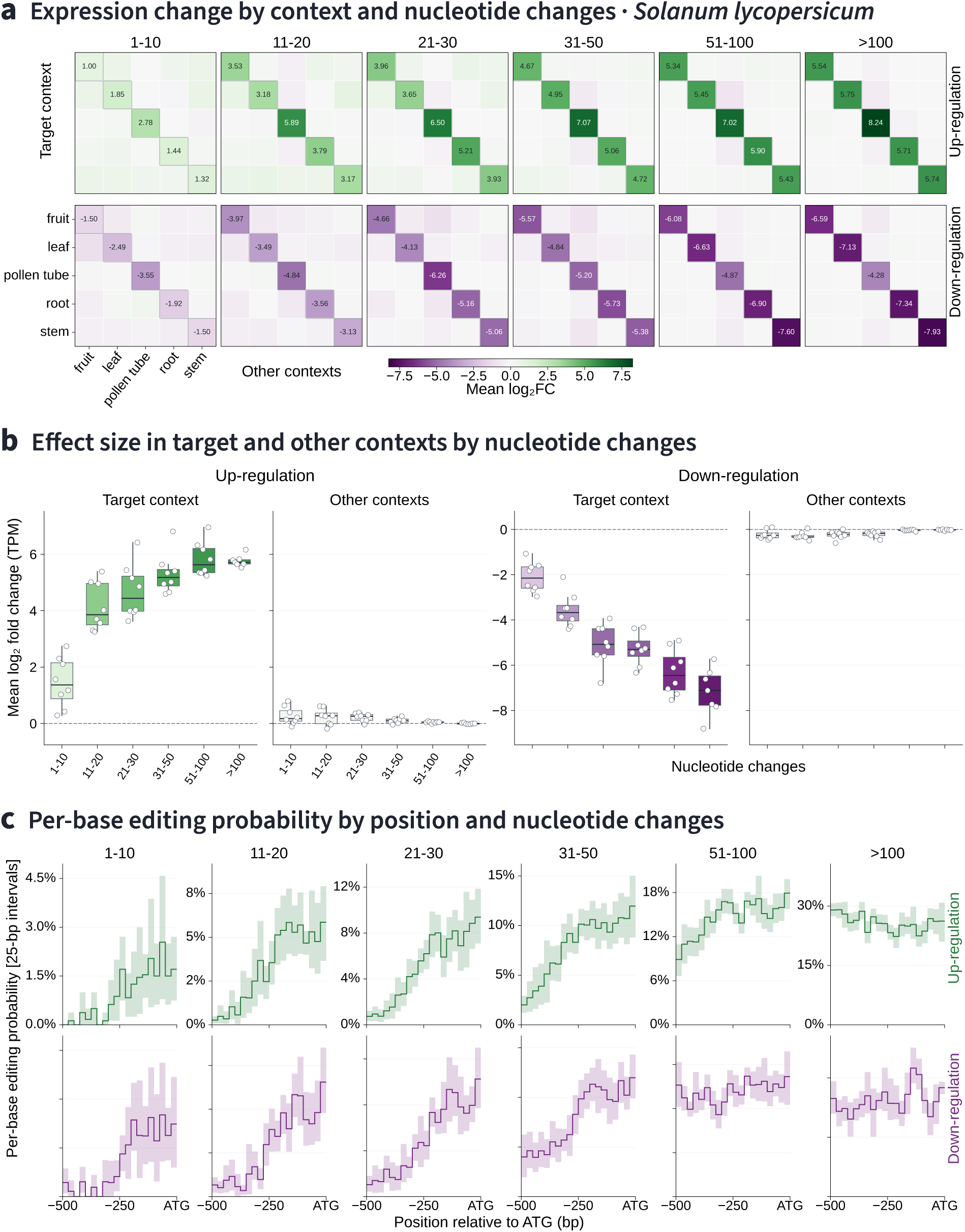
Sparse sequence edits produce context-selective predicted expression changes in tomato. Designs were generated for eight genes across fruit, leaf, pollen tube, root and stem using the workflow in Fig. 6. **a**, Mean predicted log2 fold changes, grouped by realised substitution count (1–10, 11–20, 21–30, 31–50, 51–100 and more than 100) and requested direction of regulation. Rows denote target contexts and columns denote the contexts in which expression is evaluated; labelled diagonal entries show target-context responses and off-diagonal entries show responses in the other contexts. **b**, Gene-level distributions of target responses and signed median responses across the four non-target contexts. Each point represents one gene after averaging across its available target contexts within a substitution-count bin. Boxes show the interquartile range, centre lines the median and whiskers extend to 1.5 interquartile ranges; dashed lines mark zero change. **c**, Mean per-base editing frequency in 25-bp intervals from −500 to −1 relative to the A of ATG, shown separately for up-regulation (green) and down-regulation (purple). Shading shows pointwise 95% percentile intervals from 5,000 whole-gene bootstrap resamples. Expression conversion, equal-gene weighting and position analysis follow Fig. 7 and are described in Methods.

## References

1. Rodríguez-Leal, D. et al. Engineering quantitative trait variation for crop improvement by genome editing. Cell 171, 470–480.e8 (2017). 10.1016/j.cell.2017.08.030

2. Shi, J. et al. ARGOS8 variants generated by CRISPR–Cas9 improve maize grain yield under field drought stress conditions. Plant Biotechnology Journal 15, 207–216 (2017). 10.1111/pbi.12603

3. Song, X. et al. Targeting a gene regulatory element enhances rice grain yield by decoupling panicle number and size. Nature Biotechnology 40, 1403–1411 (2022). 10.1038/s41587-022-01281-7

4. Lou, H. et al. Engineering source-sink relations by prime editing confers heat-stress resilience in tomato and rice. Cell 188, 530–549.e20 (2025). 10.1016/j.cell.2024.11.005

5. Novakovsky, G. et al. Obtaining genetics insights from deep learning via explainable artificial intelligence. Nature Reviews Genetics 24, 125–137 (2023). 10.1038/s41576-022-00532-2

6. Barbadilla-Martínez, L., Klaassen, N., van Steensel, B. & de Ridder, J. Predicting gene expression from DNA sequence using deep learning models. Nature Reviews Genetics 26, 666–680 (2025). 10.1038/s41576-025-00841-2

7. Avsec, Ž., et al. Effective gene expression prediction from sequence by integrating long-range interactions. Nature Methods 18, 1196–1203 (2021). 10.1038/s41592-021-01252-x

8. Linder, J. et al. Predicting RNA-seq coverage from DNA sequence as a unifying model of gene regulation. Nature Genetics 57, 949–961 (2025). 10.1038/s41588-024-02053-6

9. Avsec, Ž., et al. Advancing regulatory variant effect prediction with AlphaGenome. Nature 649, 1206–1218 (2026). 10.1038/s41586-025-10014-0

10. Mendoza-Revilla, J. et al. A foundational large language model for edible plant genomes. Communications Biology 7, 835 (2024). 10.1038/s42003-024-06465-2

11. Zhai, J. et al. PlantCAD2: A DNA foundation model for interpreting genomes across flowering plants. Cell Genomics, 101329 (2026). 10.1016/j.xgen.2026.101329

12. Boshar, S. et al. A foundational model for joint sequence-function multi-species modeling at scale for long-range genomic prediction. bioRxiv (2025). 10.64898/2025.12.22.695963

13. Barozet, A. et al. BOTANIC-1: a series of long-context plant genomic foundation models in the agentic era. bioRxiv (2026). 10.64898/2026.09.04.749355

14. O’Malley, R. C. et al. Cistrome and epicistrome features shape the regulatory DNA landscape. Cell 165, 1280–1292 (2016). 10.1016/j.cell.2016.04.038

15. Jores, T. et al. Synthetic promoter designs enabled by a comprehensive analysis of plant core promoters. Nature Plants 7, 842–855 (2021). 10.1038/s41477-021-00932-y

16. Jores, T. et al. Plant enhancers exhibit both cooperative and additive interactions among their functional elements. The Plant Cell 36, 2570–2586 (2024). 10.1093/plcell/koae088

17. Fu, X. et al. A foundation model of transcription across human cell types. Nature (2025). 10.1038/s41586-024-08391-z

18. Aksu, E. D. & Vingron, M. Context-aware sequence-to-function model of human gene regulation. Nature Communications 17, 6200 (2026). 10.1038/s41467-026-75527-2

19. Gao, Z. et al. RegFM: an interpretable context-aware foundation model for human transcriptional regulation. bioRxiv (2026). 10.64898/2026.08.17.744355

20. Washburn, J. D. et al. Evolutionarily informed deep learning methods for predicting relative transcript abundance from DNA sequence. Proceedings of the National Academy of Sciences USA 116, 5542–5549 (2019). 10.1073/pnas.1814551116

21. Zhai, J. et al. Cross-species modeling of plant genomes at single-nucleotide resolution using a pretrained DNA language model. Proceedings of the National Academy of Sciences USA 122, e2421738122 (2025). 10.1073/pnas.2421738122

22. Kaplan, J. et al. Scaling laws for neural language models. arXiv 2001.08361 (2020). https://arxiv.org/abs/2001.08361

23. Hoffmann, J. et al. Training compute-optimal large language models. Advances in Neural Information Processing Systems 35, 30016–30030 (2022). 10.52202/068431-2176

24. Sorscher, B. et al. Beyond neural scaling laws: beating power law scaling via data pruning. Advances in Neural Information Processing Systems 35, 19523–19536 (2022). 10.52202/068431-1419

25. de Boer, C. G. et al. Deciphering eukaryotic gene-regulatory logic with 100 million random promoters. Nature Biotechnology 38, 56–65 (2020). 10.1038/s41587-019-0315-8

26. de Boer, C. G. & Taipale, J. Hold out the genome: a roadmap to solving the cis-regulatory code. Nature 625, 41–50 (2024). 10.1038/s41586-023-06661-w

27. Vaishnav, E. D. et al. The evolution, evolvability and engineering of gene regulatory DNA. Nature 603, 455–463 (2022). 10.1038/s41586-022-04506-6

28. Gosai, S. J. et al. Machine-guided design of cell-type-targeting cis-regulatory elements. Nature 634, 1211–1220 (2024). 10.1038/s41586-024-08070-z

29. de Almeida, B. P. et al. Targeted design of synthetic enhancers for selected tissues in the Drosophila embryo. Nature (2024). 10.1038/s41586-023-06905-9

30. Anzalone, A. V. et al. Search-and-replace genome editing without double-strand breaks or donor DNA. Nature 576, 149–157 (2019). 10.1038/s41586-019-1711-4

31. Lin, Q. et al. Prime genome editing in rice and wheat. Nature Biotechnology 38, 582–585 (2020). 10.1038/s41587-020-0455-x

32. Cardi, T. & Vitale, A. Evolution and implementation of EU regulation on new genomic techniques. Nature Plants 12(9), 1650–1655 (2026). 10.1038/s41477-026-02402-9

33. European Parliament and Council of the European Union. Regulation (EU) 2026/1388 of 17 June 2026 on plants obtained by certain new genomic techniques and their products, and amending Regulation (EU) 2017/625. Official Journal of the European Union (2026). https://eur-lex.europa.eu/eli/reg/2026/1388/oj/eng

34. Liu, J. et al. PlantExp: a platform for exploration of gene expression and alternative splicing based on public plant RNA-seq samples. Nucleic Acids Research 51, D1483–D1491 (2023). 10.1093/nar/gkac917

35. Jin, J. et al. PlantTFDB 4.0: toward a central hub for transcription factors and regulatory interactions in plants. Nucleic Acids Research 45, D1040–D1045 (2017). 10.1093/nar/gkw982

36. Tian, F. et al. PlantRegMap: charting functional regulatory maps in plants. Nucleic Acids Research 48, D1104–D1113 (2020). 10.1093/nar/gkz1020

37. Schreiber, J. et al. Programmatic design and editing of cis-regulatory elements. bioRxiv (2025). 10.1101/2025.04.22.650035

38. Avsec, Ž., et al. Base-resolution models of transcription-factor binding reveal soft motif syntax. Nature Genetics 53, 354–366 (2021). 10.1038/s41588-021-00782-6

39. Jolma, A. et al. DNA-dependent formation of transcription factor pairs alters their binding specificity. Nature 527, 384–388 (2015). 10.1038/nature15518

40. Soufi, A. et al. Pioneer transcription factors target partial DNA motifs on nucleosomes to initiate reprogramming. Cell 161, 555–568 (2015). 10.1016/j.cell.2015.03.017

41. Bahri, Y. et al. Explaining neural scaling laws. Proceedings of the National Academy of Sciences USA 121, e2311878121 (2024). 10.1073/pnas.2311878121

42. Jevtic, S. et al. A scalable method for modulating plant gene expression using a multispecies genomic model and protoplast-based massively parallel reporter assay. bioRxiv (2024). 10.1101/2024.12.05.626999

43. Brookes, D., Park, H. & Listgarten, J. Conditioning by adaptive sampling for robust design. Proceedings of Machine Learning Research 97, 773–782 (2019). https://proceedings.mlr.press/v97/brookes19a.html

44. Fannjiang, C. et al. Conformal prediction under feedback covariate shift for biomolecular design. Proceedings of the National Academy of Sciences USA 119, e2204569119 (2022). 10.1073/pnas.2204569119

45. Yu, H., Liu, C. & Dixon, R. A. A gene-editing/complementation strategy for tissue-specific lignin reduction while preserving biomass yield. Biotechnology for Biofuels 14, 175 (2021). 10.1186/s13068-021-02026-5

46. Jiang, Y.-Y. et al. Prime editing efficiently generates W542L and S621I double mutations in two ALS genes in maize. Genome Biology 21, 257 (2020). 10.1186/s13059-020-02170-5

47. Vu, T. V. et al. Optimized dicot prime editing enables heritable desired edits in tomato and Arabidopsis. Nature Plants 10, 1502–1513 (2024). 10.1038/s41477-024-01786-w

48. Yates, A. D., et al. Ensembl 2026. Nucleic Acids Research 54, D1053–D1060 (2026). 10.1093/nar/gkaf1239

49. McIntosh-Smith, S., Alam, S. & Woods, C. Isambard-AI: a leadership-class supercomputer optimised specifically for Artificial Intelligence. In Cray User Group (CUG 2024) (ACM, 2024). 10.1145/3725789.3725794

